# Genome archaeology of two laboratory *Salmonella enterica enterica* sv Typhimurium

**DOI:** 10.1101/2021.04.07.438813

**Authors:** Julie Zaworski, Oyut Dagva, Anthony W Kingston, Alexey Fomenkov, Richard D. Morgan, Lionello Bossi, Elisabeth A. Raleigh

## Abstract

The Salmonella research community has used strains and bacteriophages over decades, exchanging useful new isolates among laboratories for study of cell surface antigens, metabolic pathways and restriction-modification studies. Here we present the sequences of two laboratory *Salmonella* strains (STK005, an isolate of LB5000; and its descendant ER3625). In the ancestry of LB5000, segments of ~15 and ~42 kb were introduced from *Salmonella enterica* sv Abony 103 into *Salmonella enterica* sv Typhimurium LT2, forming strain SD14; this strain is thus a hybrid of *S. enterica* isolates. Strains in the SD14 lineage were used to define flagellar antigens from the 1950s to the 1970s, and to define three restriction-modification systems from the 1960s to the 1980s. LB5000 was also used as host in phage typing systems used by epidemiologists. In the age of cheaper and easier sequencing, this resource will provide access to the sequence that underlies the extensive literature.

## INTRODUCTION

Bacteria, phages and conjugal plasmids are in a constant arms race to develop respectively defense against DNA invasion and methods of evading defenses. Bacteria express variable surface features to thwart entry (e. g. flagellae, outer membrane proteins, O-antigen sugar chains) and internal limitation mechanisms to abort establishment (e. g. restriction systems and abortive-infection suicide factors) (Doron *et al.* 2018; Azam and Tanji 2019). Phages and plasmids may defeat host restriction-modification (RM) systems with antirestriction proteins, which frequently block type I DNA restriction-modification enzymes (e. g. (Zavilgelsky and Rastorguev 2009; Serfiotis-Mitsa *et al.* 2010; Balabanov *et al.* 2012; Roberts *et al.* 2012; Piya *et al.* 2017; Gonzalez-Montes *et al.* 2020). Low-specificity DNA modification enzymes expressed only for the entering molecule are also found in prophages (Murray *et al.* 2018) and plasmids (Fomenkov *et al.* 2020).

Classical studies of horizontal exchange processes within and between genera frequently employed strains in the ancestry of LB5000, particularly for study of cell surface antigens and restriction systems. Early hybrid serovar ancestors were constructed by transduction from *S. enterica* sv Abony into *S. enterica* sv Typhimurium. These studies defined flagellar antigen types, the phenomenon of flagellar phase variation (Joys and Stocker 1965; Joys and Stocker 1969) and the use of O-antigen as phage receptor (Ornellas and Stocker 1974; Stocker *et al.* 1980). Three restriction systems were identified using this hybrid lineage (Bullas and Colson 1975). In LB5000 itself, the three restriction systems of strain LT2 were inactivated (Bullas and Ryu 1983), allowing use for epidemiologic purposes as a host for phages in the Anderson phage typing system (Schmieger 1999). Intergeneric conjugal transfer of chromosomal markers was studied using *Escherichia coli* K-12 derivatives and *Salmonella enterica* sv Typhimurium lysotypes LT2 and LT7, using F’ plasmids and integrated F factors (Sanderson 1987). LB5000 and relatives are particularly useful as an intermediate host in molecular genetic applications, moving constructions between strains and genera (Sanderson and Roth 1988; Segall and Roth 1989; Chaudhuri *et al.* 2009; Lerche *et al.* 2017; Yoon *et al.* 2018; Griewisch *et al.* 2020; Jones-Carson *et al.* 2020).

Genome sequences are available for LT2 (Mcclelland *et al.* 2001) and LT7 (Zaworski *et al.* 2021), but none within this hybrid lineage.

Here we present annotated sequences of two *S. enterica* sv Typhimurium variant genomes: STK005 (our isolate of LB5000, from Anca Segall’s G251), and ER3625, a derivative of STK005 built in our lab for studies of HGT events. The LB5000 genome carries scars from genetic steps in its descent from LT2, including the replacement of two flagellum-encoding segments of the LT2 genome by segments of *S. enterica* sv Abony, with concomitant loss of prophage Fels2. Recombination events involving two prophages document intralineage allele shuffling. Mutations linked to the amino acid requirement of the strain were identified as well.

## MATERIAL AND METHODS

### *Salmonella* growth

For each strain, a single colony was grown in RB (10 g soy peptone, 5 g yeast extract, 5 g NaCl per liter) with the appropriate antibiotics (kanamycin 40 μg/ml, streptomycin 100 μg/ml) overnight at 37°C with 250 rpm agitation.

### DNA preparation, sequencing and de novo assembly

#### DNA extraction

STK005 and ER3625 were prepared for PacBio RSII or Sequel sequencing. gDNA was extracted with the Monarch Genomic DNA purification kit (#T3010 New England Biolabs; Ipswich, MA, USA). For Oxford Nanopore sequencing of STK005, gDNA was extracted from 1 ml of overnight culture with the kit Monarch HMW DNA Extraction kit (T#3060 New England Biolabs; Ipswich, MA, USA) following the manufacturer instructions for High Molecular Weight DNA extraction from gram-negative bacteria. After the extraction, the HMW DNA was homogenized following by heat and pipetting before QC. A clean peak > 60 kb was found on analysis with the Genomic DNA ScreenTape® (Agilent Technology, Santa Clara, CA).

#### Library preparation

For the PacBio RSII sequencing platform, STK005 SMRTbell libraries were constructed from genomic DNA samples sheared to between 10 and 20 kb using the G-tubes protocol (Covaris; Woburn, MA, USA), end repaired, and ligated to PacBio hairpin adapters. Incompletely formed SMRTbell templates and linear DNAs were digested with a combination of Exonuclease III and Exonuclease VII (New England Biolabs; Ipswich, MA, USA). DNA qualification and quantification were performed using the Qubit fluorimeter (Invitrogen, Eugene, OR) and 2100 Bioanalyzer (Agilent Technology, Santa Clara, CA). The SMRTbell library was prepared according to PacBio sample preparation protocol sequenced with C4-P6 chemistry with a 300 min collection time. A total of 751,460 polymerase reads with N50 length of 17,626 bp was obtained (approximately 360X coverage) before filtering.

ER3625 libraries were sequenced using the PacBio Sequel I platform. Briefly, SMRTbell libraries were constructed from genomic DNA samples following the PacBio protocol for Sequel using the kit 100-938-900. DNA qualification and quantification were performed using the Qubit fluorimeter (Invitrogen, Eugene, OR) and 2100 Bioanalyzer (Agilent Technology, Santa Clara, CA). The resulting library, of average size 11,640 bp, was prepared for binding following the PacBio guidelines generated by SMRT Link and run on a Sequel I machine.

The STK005 Nanopore library was prepared with 1.7 μg of HMW DNA following the Genomic DNA by ligation (SQK-LSK109 protocol) from Oxford Nanopore. The adapter ligation was performed for 1 h instead of 10 min. The flow cell was primed and loaded with ~100 ng of library following the same documentation. After a 21h run on a second-use FLO-MIN106 flow cell with the GridIONxs apparatus, 449K reads were obtained, with 27.39 kb read length basecalled N50. The basecalling was performed on the fly with the high-accuracy basecalling model from Guppy (version 3.2.10)

#### Sequence assembly

The STK005 (PacBio RSII reads) were assembled using RS_HGAP3 program with parameters: minimum subread length 5,000 bp, minimum polymerase quality 0.8, Minimum polymerase read length 10,000 bp, 5 Mbp genome size and other parameters as default settings. The three resulting contigs, of size 4,802,499 bp, 103,465 bp and 8,138 bp, were circularized with Circlator (Hunt *et al.* 2015) with CCS generated with parameters minimum full passes 3, minimum predicted accuracy 95, minimum read length of insert 1000 bp and the other parameters as default. The smallest contig was a subset of the chromosome contig.

PacBio polishing for STK005 was done by running the RS_Reseq program twice, with Minimum subread length 500, minimum polymerase read quality 75 and then 80, and 1500 bp for Minimum polymerase read length. The final assembly is composed of a 4,796,208 bp chromosome and a 93,938 bp plasmid.

The STK005 Nanopore reads were corrected and filtered using Canu (Koren *et al.* 2017), the output corrected reads were then trimmed and assembled with Canu (version 1.4). A first correction step was run with parameters minReadLength = 10000, stopOnReadQuality=false and other as default. Then trimming and assembly were run with genomeSize= 5m and other parameters as default. Two contigs of length 162,546 and 4,863,042 bp corresponding to the plasmid and chromosome.

In parallel, the reads were mapped on the PacBio de novo assembly using minimap2 (version 2.17) with default parameters (Li 2018). In both cases, no inversion in between the Gifsy prophages was observed.

PacBio Sequel assembly of ER3625 used the RS_HGAP4 program with parameters: 5 Mb expected genome size, minimum mapped length 5 kb, minimum mapped concordance 70 and other parameters as default settings. The 2 contigs obtained were circularized with Circlator (version 1.5.5). Polishing was done by running the RS_Reseq program with minimum mapped length 500, yielded 19 corrections. The final assembly is composed of a 4,796,355 bp chromosome (coverage 541x) and a 93,939 bp plasmid (2,247x coverage).

#### Modification motifs

DNA motifs and degree of modification were generated using InterPulse Duration (IPD) Ratios analysed with RS_Modification_and_Motif_Analysis from PacBio as in (Flusberg *et al.* 2010; Clark *et al.* 2012).

#### Genome annotation

Genbank deposits STK005 (chromosome CP067397, plasmid CP067398) and ER3625 (chromosome CP067091 and plasmid CP067092) genomes were annotated with the Genbank PGAP pipeline (Tatusova *et al.* 2016; Haft *et al.* 2018). In-process analysis employed annotations in Geneious (2018 version in steps: Menu Align and Assemble; Align Whole Genomes; Tab Alignment View; Save as editable copy; Menu Annotate and Predict; Annotate from Database) using LT2 [NC_003197.2] and Abony [NZ_CP007534.1] and segment extractions from them as reference genomes in a local database folder.

### Data availability

Supplementary material is available at figshare DOI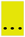 Table S1 “Pedigree LB5000 STK005” traces the descent of LB5000 and some relatives through the literature since 1959; with wild ancestor, donor strain, recipient strain, renaming, construction reference and mode of construction. File S1 “STK005 variants” contains 5 Excel spreadsheets documenting positions at which STK005 sequence differs from the genome sequences representing two ancestral components. For three genome segments derived from LT2, variant nucleotide positions of LT2 (NC_003197.2) are listed, with consequences for amino acid changes; for two short segments derived *Salmonella enterica* serovar Abony SW803 (H1, 15.4 kb and H2, 41.7 kb), all annotated loci are listed as well, since few positions vary from the Abony reference. The reference segments H1 and H2 were extracted from *Salmonella enterica subsp. enterica* serovar Abony str. 0014 - NZ_CP007534.1: for transduction 1 of Table S1, H1--nt 3956473-3942278 (reversed); for transduction 2, H2--nt 4713943-4725447(catenated)1-18493 (the segment crosses the numerical origin, so is catenated). File S2 “Analysis of 11 kb high-similarity region in the Gifsy prophages” describes and displays alignments (referred to as “Embedded figs“) that resolve allele configurations in Gifsy prophages. File S3 “STK005 genotype inferences from sequence” describes the recorded genotype; explains assignment of flagellar genotype; describes the protocol used to generate the variant list found in File S1; lists protein changes that account for the genotype; and points out some additional possible changes. File S4 “PCR survey for 42bp insertion in isolates of *S. enterica* sv Typhimurium str LT2” demonstrates the insertion is present in Gifsy-1 and conserved in most LT2 strains.

All the sequencing data can be found in the BioProject PRJNA605961. The genome sequences of STK005 and ER3625 are available from GenBank with the accession numbers (CP067397-CP067398) and (CP067091-CP067092), respectively. All the raw data are available under the SRA ID SRR13390529, (SRR13404095, SRR13404094, SRR13404093, SRR13404092, SRR13404091) and SRR13787361 for ER3625, STK005 RSII and STK005 Nanopore respectively. The methylation data are available as supplementary files on NCBI under the IDs SUPPF_0000003906 and SUPPF_0000003907. Both strains are available from New England Biolabs.

## RESULTS AND DISCUSSION

### Strain engineering scars: *Salmonella enterica* subsp. *enterica* serovar Abony DNA in LB5000

The engineering history of LB5000 (Figure 1) includes two transductions from *Salmonella* serovar Abony strain 803 (Spicer and Datta 1959) (see details and references in Table S1). These two transductions left scars in the form of high-divergence segments in the genome surrounding the flagellar loci H1 and H2 (Figure 2A). In *Salmonella enterica*, both H1 and H2 flagellar antigens are highly variable in wild lineages in the population; in addition, each individual lineage is capable of switching from expressing its H1 antigen to its H2 or the reverse, a phenomenon known as phase variation (Simon *et al.* 1980). This case of phase variation involves reversible site-specific rearrangement of regulatory signals that enable expression from only one of two distant loci. In LB5000, both of these loci derive from *S. enterica* sv Abony.

**Figure 1:**
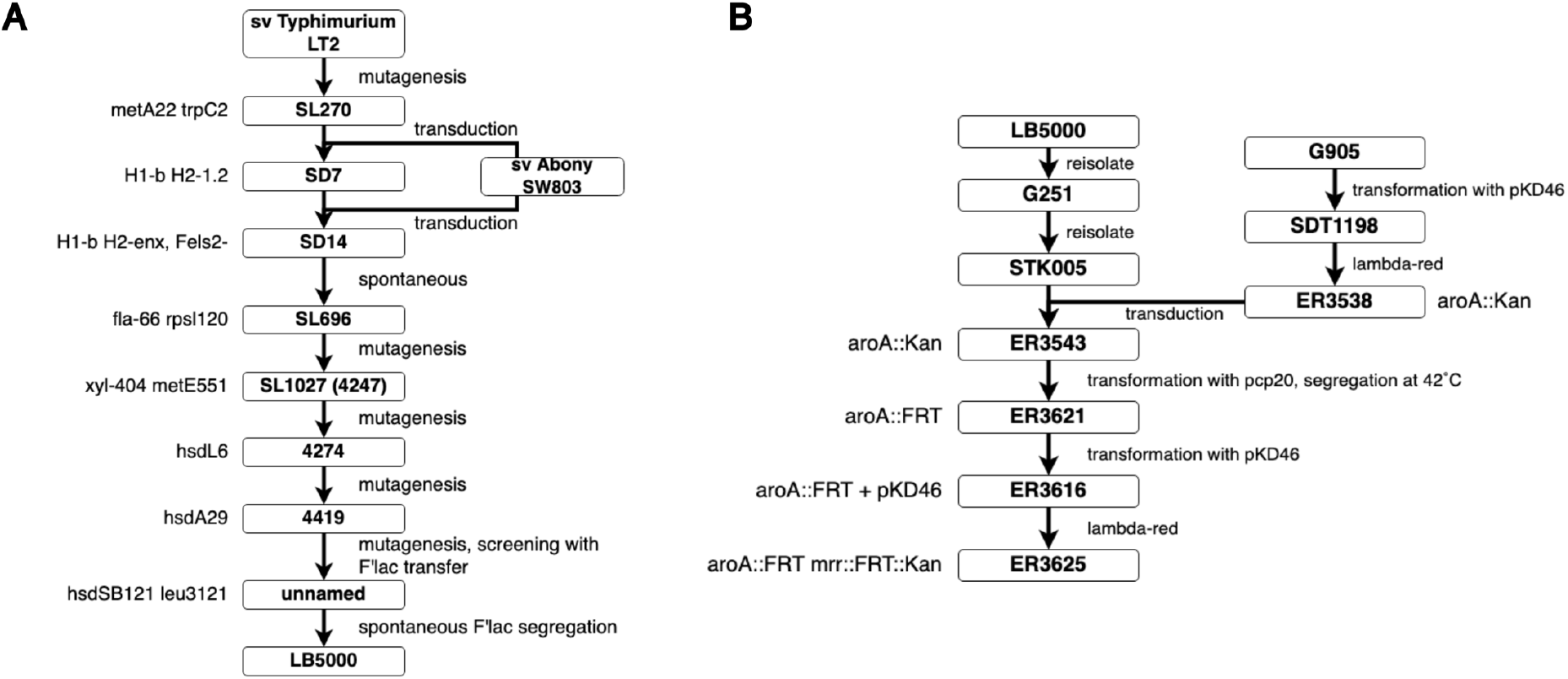
Strain pedigree charts. A. LB5000 B. ER3625. Strain names are boxed. To the right of each construction step (arrows) are the methods used; to the left of each box is the genotype designation assigned as a result of the step. In some cases, a genotype marker was discovered during later investigation. The different strain modifications were done by mutagenesis (random by use of NG), transduction (use of phages) or transformation (plasmids with induction). For more details and references on these steps, refer to Table S1.

**Figure 2:**
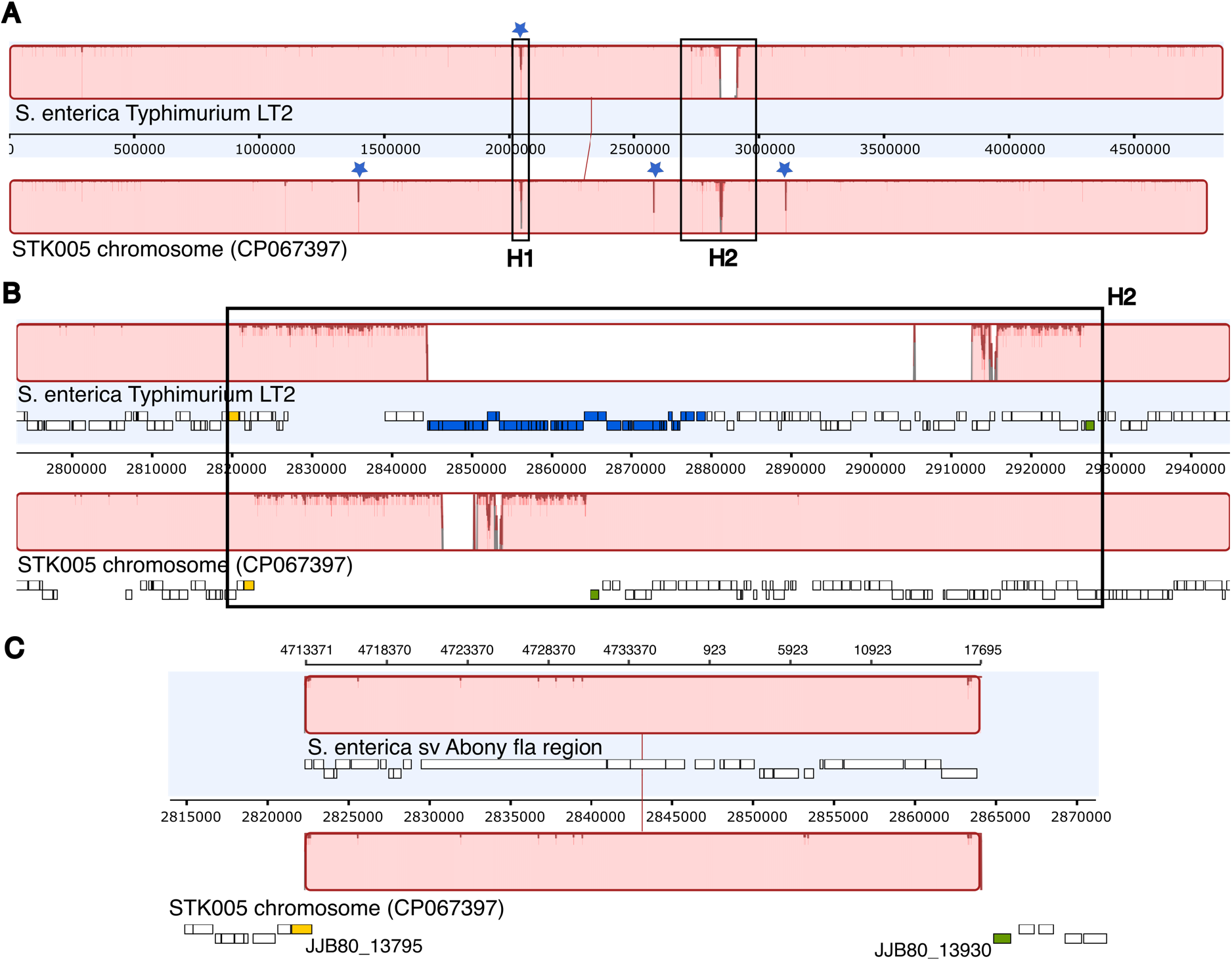
Recombination scars. Panel A is a Mauve alignment of the STK005 and LT2 chromosomes. Blue stars represent IS sequences not shared between the sequences. The flagellar regions H1 and H2 are boxed. Light pink color is for high similarity areas; the white area is a missing segment; and dark red lines dip where mismatches occur. Panel B is a close-up of the H2 region. Panel C is a Mauve alignment of S. Abony H2 (fla) region with STK005. In panel B and C, the yellow and green rectangles below the alignment are landmarks, respectively gene STM2679 (JJB80-13795) and STM2780 (JJB80-13930).

#### H1, phase 1 flagellin region transduction patch

The H1 phenotype depends on the allelic state of the gene *fliC*, within the H1 box of Figure 2A. This patch is the result of a first transduction mediated by P22: recipient strain LT2 SL270 replaced the H1-i 1.2 allele of LT2 with H1-b from *S.* Abony SW803 (Spicer and Datta 1959; Joys and Stocker 1969) (see Table S1). Allele H1-b was bequeathed to LB5000 ancestor SD7. The highly divergent interval (LT2 locus_ID STM1946 (*uvrC*) to STM1961 (*fliD*)) includes two unshared IS insertions (IS3 in STK005, IS200 in LT2) and encompasses the *fliC* flagellin gene. In contrast, this region in LB5000 (JJB81_10090 to JJB81_10010; nt 2037077 -> 2052486) aligns with only three mismatches to *S.* Abony (SEEA0014_RS19195 to SEEA0014_RS19270; nt 3941779 -> 3957191); see below and File S1.

#### H2, phase 2 flagellin region transduction patch

The H2 phenotype is determined by the allelic state of the gene *fljB*, a *fliC* homolog. When the adjacent regulatory gene *fljA* is expressed, it mediates repression of distant H1 expression; when the local DNA segment is inverted, *fljA* and *fljB* are not expressed, and repression of distant gene *fliC* is lifted. Wild type for this LT2 isolate is H2-1.2; *S*. Abony SW803 was H2-enx. H2-enx was bequeathed to LB5000 ancestor SD14.

These genetic changes are reflected in a second divergent region of the alignment (H2 black rectangle in Figure 2A; closer view in Figure 2B). Genes present in LT2 but absent in STK005 include the Fels2 prophage (STM2694 to STM2740) and the flanking genes STM2741 to STM2769. Present but divergent are genes JJB81_13790 to JJB81_13790 when aligned with LT2 locus_ID STM2679 and STM3779. We infer that this divergent region represents genomic replacement of 106 kb of *S.* sv Typhimurium LT2 by 41 kb of *S.* sv Abony 803 during the second transduction (Figure 1, panel A). Confirmation of the inference is illustrated in Figure 2C: 41 kb of our sequence is very similar (3 insertions, 14 mismatches) to a 41 kb segment of NZ_CP007534 (sequence of *S. enterica* sv Abony strain 0014 (NCTC 6017; ATCC BAA-2162) (Timme *et al.* 2013); see File S1 sheet 4). Both flanks of the long deletion show high divergence from LT2, and the *fljB* region is 100% identical to the sequenced *S.* sv Abony genome. Prophage Fels-2 is not present in Abony serovar isolates, consistent with the present configuration.

#### *hin* gene, STM2772, JJB81_13890

The Hin protein mediates inversion of a segment including its own gene and a promoter of *fliC* (H2, phase 2) and the *fljA* (repressor of *fliC* (H1, phase 1)) (Simon *et al.* 1980). The STK005 *hin* homolog (JJB81_13890) is within the H2 transduction patch. It is similar to the Abony SEEA0014-RS00045 gene except for the 3 single nucleotide insertions; these insertions are all shared among LT2, STK005 and ER3625. In addition to the closer identity to *S. enterica* sv Abony, the *hin* gene has suffered an inversion during the descent of STK005 from LT2 (also found in ER3625). Considering the expected protein activity, this reorganization is evidence of expected function.

#### ER3625 engineering and variation from STK005

ER3625 was engineered in our laboratory to create a non-pathogenic recipient (Hoiseth and Stocker 1981) with no Mrr restriction, to pursue work on R-M systems in HGT using the system previously used in our laboratory (Kingston *et al.* 2015; Kingston *et al.* 2017). Strain engineering required several steps with a variety of methods (Figure 1B).

For future work we wish to verify that no off-target changes had occurred during lamda-Red-mediated genome engineering or accompanying transduction. The *mrr* engineered locus carries the designed sequence (*npt* flanked by FRT sites replacing the gene). The *aroA* gene was deleted, but the scar of the FRT site recombination is aberrant. This DNA segment is annotated by NCBI pipeline as JJB80_05020 (DUF1317 domain-containing protein) and JJB80_05025 (DUF1382 protein family). No SNPs were identified in genes between STK005 and ER3625 for either the chromosome or the plasmid.

### Structural complexity and intralineage prophage recombination

Gifsy-1 and Gifsy-2 are two LT2 prophages located between 2,728,977 and 2,776,819 (STM2584 to STM2636) and between 1,098,231 and 1,143,702 (STM1005 to STM1056) respectively. These two related prophages are patchily similar to each other; while overall DNA identity is 44.1%, this ~42 kb region is interspersed with highly divergent segments and unrelated insertions (Figure 3). Of concern are two long regions of high identity: a ~12 kb stretch of > 98% identity in the regions expressed early in lytic infection: recombination, regulators related to CI, CII and Cro of lambda, and replication; and an 8 kb patch of ~ 78% identity at the other end of prophages.

**Figure 3:**
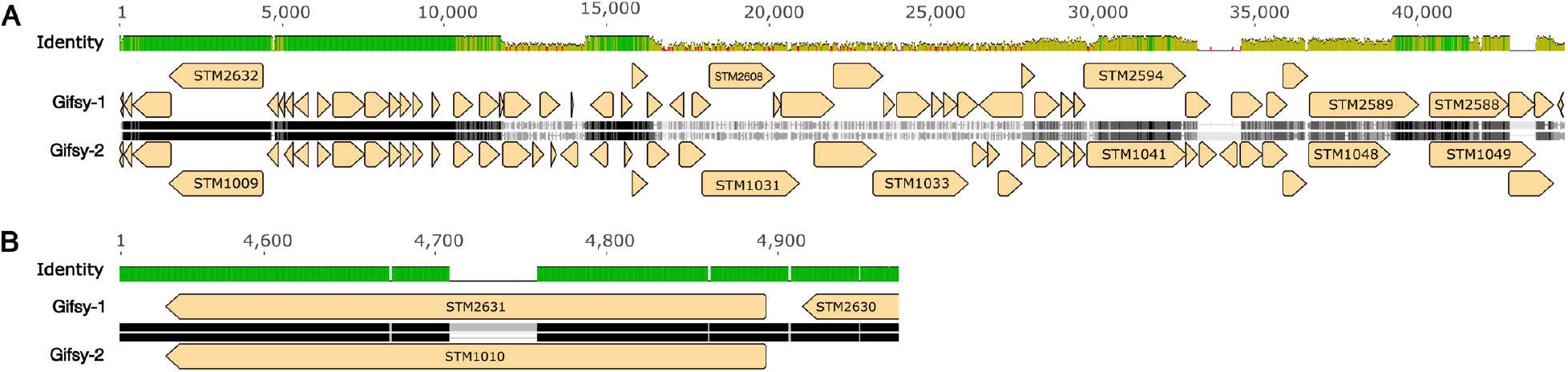
Pairwise alignment of the Gifsy-1 and Gifsy-2 prophages of LT2. Panel A: alignment of the whole prophages. Panel B: close-up of STM2631 and STM1010, revealing a 51 bp indel variation. The identity rows: green=identical bases; yellow=lower identity; red=very low local identity. The sequences Gifsy-1 and Gifsy-2 are in black, white and gray: black for identical bases, white for SNPs, gray for insertion/deletion (indel) variation (corresponding gaps shown with a thin line).

These long regions with very high identity are of concern for both technical and biological reasons. First, long repeats are an assembly challenge even with long-read technologies. Second, recombination within the host may generate new gene combinations that are then transmissible to new hosts on prophage induction. Extensive mosaicism among tailed phages is well known (Grose and Casjens 2014); and creation of a new mosaic by recombinational marker rescue from defective prophages has been noted before (De Paepe *et al.* 2014). Evidence of recombination within a laboratory lineage was found in *E. coli* ER2796 (Anton *et al.* 2015).

#### Resolution of assembly errors

De novo assembly of the STK005 genome initially yielded (artifactual) chromosome inversion. On examination of reads aligned with the two Gifsy regions, many poorly or partially mapped reads and an important number of SNPs were seen, suggesting imperfect mapping. A reassembly using only long CCS (> 5 kb; 47.7x coverage) was then created with HGAP3, then circularized with Circlator. This assembly did not show a genomic inversion relative to LT2. Further polishing to resolve SNPs used RS_reseq with CCS > 500 bp (220.9x coverage). This time, few reads were mismapped at the Gifsy regions, confirming the absence of the inversion in STK005 genome.

In parallel, Oxford Nanopore was used to generate long reads that could cover 40-45 kb regions and reinforce confidence in the structural organization of the assembly. Reads >10 kb were used to generate a de novo assembly with Canu (version 1.4). This assembly agreed with the final PacBio assembly, i.e. without inversion of the genome. Mapping of the Nanopore reads on the PacBio assembly confirmed the structure.

#### Prophage recombination

This 12 kb region of near identity codes for proteins involved in homologous recombination, as well as regulation and replication. The recombination proteins are related to those of the *E. coli* Rac prophage RecET recombination cluster; also present are homologs of Lambda Orf and Rap, which also affect recombination. RecE and RecT provide homologous recombination systems that can substitute for the host RecABCD system (Lemire *et al.* 2008; Murphy 2016). Rap (originally named NinG) facilitates recombination near linear ends via the Lambda Red pathway (Tarkowski *et al.* 2002), acting as a structure-specific nuclease (Sharples *et al.* 2004).

Three segments within this long region posed interpretation problems: assembly error or recombination during the lineage leading to STK005. Our interpretation is that the reference LT2 sequence is in error in one case, and both reciprocal recombination and gene conversion have affected this region (see Figure 3 and Files S2 and S4).

First is the appearance of a novel allele of Gifsy-1 *recET*, including a reading-frame-preserving insertion of 42 nt (14 aa) in the middle of the RecE CDS and 9 SNPs flanking it (File S2 Embedded fig2). The Gifsy-1 allele of *recET* found here appears to be ancestral: it is also found in the Gifsy-1 sequence of *S. enterica* sv Typhimurium strain 14028S (Genbank CP034479.1; File S2 Embedded fig2). (Figueroa-Bossi *et al.* 2001; Lemire *et al.* 2007). Strain 14028S was used extensively in laboratory studies due to much more virulent properties in mice (e.g. (Figueroa-Bossi *et al.* 1997)). PCR tests of 19 LT2 isolates or derivatives confirm the presence of the insertion in all of them except strains specifically cured of Gifsy-1 (File S4 Embedded fig1 and Embedded table1). A recent, independently-determined LT2 sequence (CP014051) was obtained using PacBio and Illumina technology. This deposit agrees with our sequence of STK005 across this region.

Second, there has been a reciprocal exchange of the sequence between Gifsy-1 and Gifsy-2 prophages (File S2 Embedded fig3A, 3B). This exchange includes the position of the 18-amino acid (51 bp) indel variation in CDSs encoding a DNA break and rejoin activity illustrated in Figure 3B above.

Third, gene conversion has resulted in loss of the original Gifsy-2 allele of Rap (NinG), replaced by Gifsy-1 information without change in the original Gifsy-1 (File S2 Embedded fig4A, 4B).

Finding evidence of recombination activity within the high-identity region is interesting. Though normally silent as prophages, both Gifsy-borne LT2 homologs of RecET can be activated by mutation to mediate recombination in vivo (Lemire *et al.* 2008). Action of these phage recombination systems may promote the mosaic genomic relationships among phages that has been found (Martinsohn *et al.* 2008). The two events found here are compatible with that role.

### Recorded genotype and point mutations

#### Metabolic markers

LB5000 was originally described as auxotrophic for methionine, tryptophan and leucine with the genotype *metA22 metE551 trpC2 ilv-452*. We identified mutations likely responsible for the reported phenotypes; we also found mutations in other genes involved in amino acid synthesis, see File S3.

Concerning methionine production, changes are found in *metA* (*metA22*) and *metE* (*metE551*). In *met*A (STM4182 in LT2 and JJB80_21020 in STK005), a 10 nt deletion at the protein C-terminus truncates it by 9 amino acids. A SNP in *metE* (G877A) leads to G293R protein substitution. The *trpC2* allele carries nt change G545A leading to G->D amino acid substitution. For isoleucine and leucine auxotrophies, SNPs were found in several genes. Coding changes were a G > A mutation at position 418 in *ilvC* (G140S amino acid change) and a C to T transition in *leuA* at position 424 (D142N).

Considering other amino acid pathways, *proB* and *proC* of proline synthesis have mutations respectively C > T at position 535 (P179S) and C > T at position 317 (A106V). However, our auxotrophy tests found that the strain requires methionine, valine/isoleucine and tryptophan/phenylalanine/tyrosine but can grow without addition of proline in the media (data not shown). The synthesis of proline may be mediated via *putA*, which has two silent mutations.

#### Genotype and phenotype of methyltransferases and their associates

DNA methylation motifs and level of methylation mediated by R-M systems and orphan MTases were analyzed with the PacBio RSII. At least six methylation motifs are known in ancestral strain *Salmonella enterica* serovar Typhimurium LT2 ((Roberts *et al.* 2015); REBASE Organism number 18099). Three motifs result from enterobacterial “orphan” methyltransferases (M; M.SenLT2 Dam, M.SenLT2 Dcm and M.SenLT2IV; (Roberts *et al.* 2015)), and three are associated with restriction-modification (RM) phenomena (M.SenLT2I, M.SenLT2II and M.SenLT2III). Two of these RM systems have been well characterized. Characterization began with a transfer isolate SD14 (Stocker lab strain SL1027). M.SenLT2I (known in the literature as the StyLT or StyLTI RM system (Colson and Colson 1971)) is a Type III enzyme, and M.SenLT2II (StySB or StyLTII (Fuller-Pace *et al.* 1984)) is Type I. The third system, RM.SenLT2III (StySA), is known to confer m6A modification (Hattman *et al.* 1976) but its protection/restriction mechanism of action remains unclear.

The LB5000 RM genotype is listed as *hsdL6 hsdSA29 hsdSB121* for StyLT, StySA and StySB. We identified appropriate mutations consistent with the R^−^M^+^ phenotype (Table 1 for M activity; File S3 for genotype discussion). For StyLT and StySB, sense mutations leading to amino acid changes were identified in the endonuclease genes, as well as silent mutations. Two G->A changes (position 393 and 2,749 of STM0358) are present in the *res* gene of StyLT, the second one leading to a D->N amino acid change in the C-terminal domain. For StySB, two G->A changes (position 2,769 and 2,473 of STM4526) are present in *hsdR*, one resulting in an R->C amino acid change in the C-terminal domain.

**Table 1.**
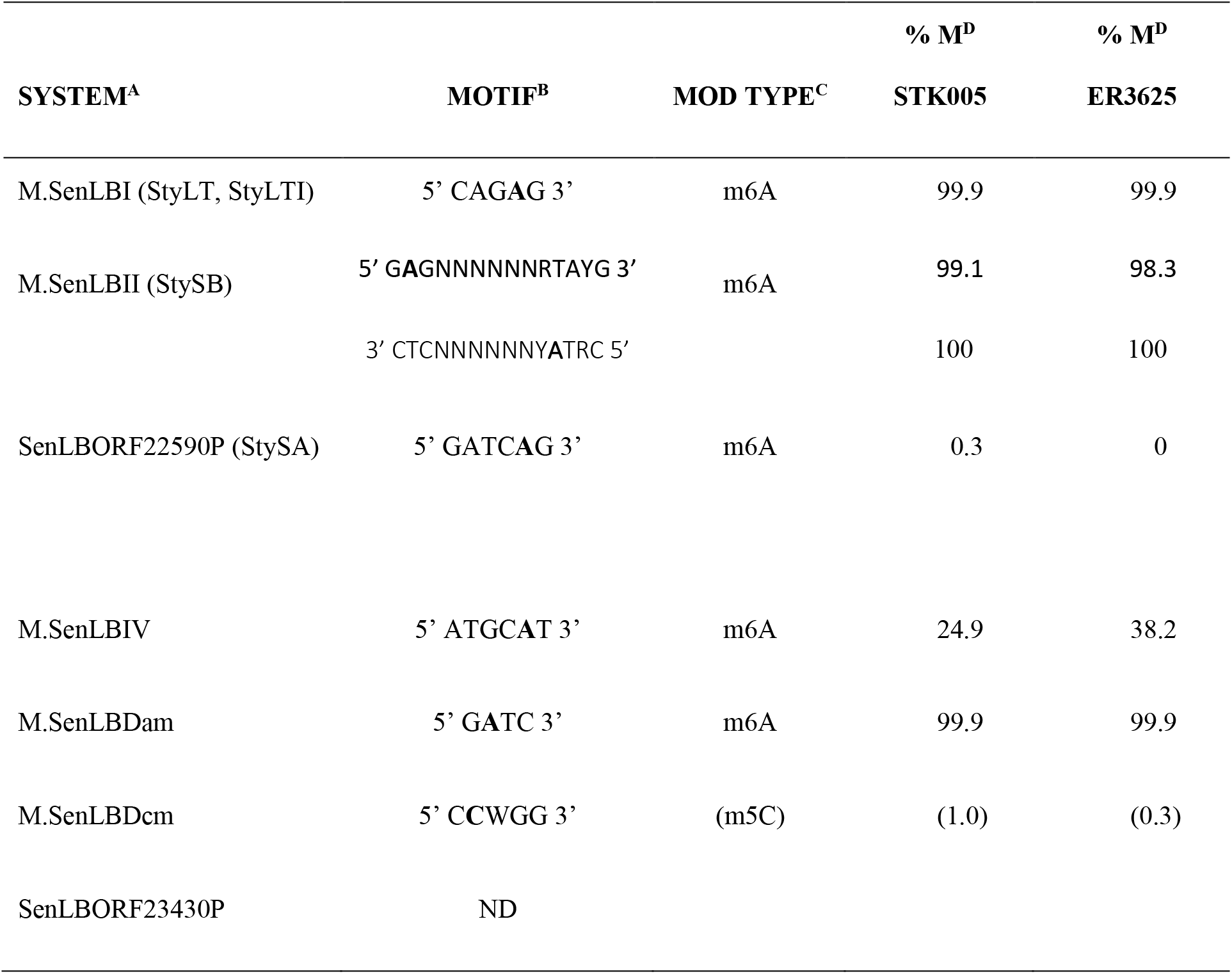
DNA modification systems, motifs and degree of modification.

The LB5000 StySA R^−^M^+^ phenotype was reported to be unstable (Tsai *et al.* 1989), and we found our STK005 isolate to be phenotypically StySA R^−^M^−^. The genetic position of StySA was mapped close to and counterclockwise of the *hsdSB* locus (Colson and Van Pel 1974); we identified a methyltransferase homolog (JJB81_22590) in an appropriate location, but found no coding change relative to STM4495. The genetic requirements for action of this system are under study.

In Table 1, we also find partial methylation for the conserved orphan methyltransferase M.SenLT2IV, as previously observed in three *Salmonella* species (Pirone-Davies *et al.* 2015). Methylation of m5C by M.Dcm was not determined here, due to technical limitations of Pacific Bioscience long-read sequencing in detection of m5C (Flusberg *et al.* 2010).

## ACKNOWLEGDGMENTS

We thank Stanley Maloy and Anca Segall (San Diego State University) for *Salmonella* strains, phages and advice and training in their use. We thank Chloé Baum et Giron Koetsier for technical assistance with the Oxford Nanopore sequencing and high molecular weight DNA extraction.

New England Biolabs provided support for JZ, OD, TK, AF, RM and EAR. The authors declare no conflicts of interest.

