## Supplemental Table S1 for "Genome archaeology of two laboratory *Salmonella enterica enterica* sv Typhimurium"

**Table S1 Pedigree, genetic steps and markers brought into LB5000 or strains used for phage propagation**

| Wild ancestor | Relevant marker in LB5000 | Strain name in reference | Reference | Original name (other names) | Other reference | Recipient or strain mutagenized | Donor if gene transfer | construction, comment Reference | Mode of construction, phenotype comment |
|---|---|---|---|---|---|---|---|---|---|
| *Salmonella enterica* sv Abony (Abony) | §H1-b H2-e,n,x, Fels2- | SW803 | (Spicer and Datta 1959) | *S. abony* 103 | (original) EDWARDS, P.R. and BRUNER, D. W. (1942). Serological identification of Salmonella cultures. Circ. K y Agric. Exp. Stn, no. 54. | | | | |
| *Salmonella enterica* sv Typhimurium LT2 (LT2) | Framework | LT2 | (Spicer and Datta 1959) | | (original) (Zinder and Lederberg 1952) | LT2 | | | |
| LT2 | *metA22 trpC2* | SL270 | (Spicer and Datta 1959) | | | LT2 | | | mutagenesis-->metA- trpB- |
| LT2 x Abony | b1.2 | SD7 | (Spicer and Datta 1959) | | | SL270 | SW803 | (Joys and Stocker 1969) | Transduction A-->i 1.2->b-1.2; fla+ phase I non-I with HI of Abony, antigen b. *nml* was cotransduced. (N-methyllysine |

| Wild ancestor | Relevant marker in LB5000 | Strain name in reference | Reference | Original name (other names) | Other reference | Recipient or strain mutagenized | Donor if gene transfer | construction, comment Reference | Mode of construction, phenotype comment |
|---|---|---|---|---|---|---|---|---|---|
| | | | | | | | | | decoration on flagellin) |
| LT2 x Abony | H1-b H2-e,n,x, Fels2- | SD14 | (Spicer and Datta 1959) | | (additional genotype information in (Joys and Stocker 1969; Wilkinson et al. 1972)) | SD7 | SW803 | (Spicer and Datta 1959) | Transduction B 1.2-->b-enx Fels2- |
| LT2 x Abony | *flaA66 rpsL120* unmapped additional mutation affecting StrR | SL696 | (Joys and Stocker 1965) and Smith and Stocker unpub cited in Wilkinson et al (Wilkinson et al. 1972) | (other name SGSC1649) | (additional genotype information in (Wilkinson et al. 1972)) | SD14 | | | spontaneous mutation--> Fla- high StrR Rfa+ |
| LT2 x Abony | *xyl-404 metE551* | SL1027 | (Wilkinson et al. 1972) | | | SL696 | | (Kuo and Stocker 1970) | mutagenesis-->xyl- metE- |
| LT2 x Abony | interlab transfer, new name | 4247 | (Bullas and Colson 1975) | SL1027 | (Wilkinson et al. 1972) | | | | |
| LT2 x Abony | *hsdL6* | 4274 | (Bullas and Colson 1975) | | | 4247 | | | NG mutagenesis R-M+ LT |
| LT2 x Abony | sister lineage to LB5000 (allows growth of | 4278 | (Bullas and Colson 1975) | | | 4247 | | | NG mutagenesis R-M- LT |

| Wild ancestor | Relevant marker in LB5000 | Strain name in reference | Reference | Original name (other names) | Other reference | Recipient or strain mutagenized | Donor if gene transfer | construction, comment Reference | Mode of construction, phenotype comment |
|---|---|---|---|---|---|---|---|---|---|
| | unmodified phage) | | | | | | | | |
| LT2 x Abony | sister lineage to LB5000 (allows growth of unmodified phage) | 4414† | (Bullas and Colson 1975) | | | 4274* | | | NG mutagenesis R-M- SA |
| LT2 x Abony | *hsdSA29* | 4419† | (Bullas and Colson 1975) | (other names CL4419, SL1654, SGSC261) | | 4274* | | (Bullas and Colson 1975) | NG mutagenesis R-M+ SA |
| LT2 x Abony | sister lineage to LB5000 (allow P1 transduction) | 4526 | (Bullas and Colson 1975) | | | 4419 | | | phage Felix-O resistance selection-->Gal- P1S |
| LT2 x Abony | *hsdSB121* | unnamed | (Bullas and Ryu 1983) | | | 4419 | | | NG mutagenesis, screening with F' lac for increased Lac+ transfer; allele number culled from SGSC listings of descendants of LB5000 |
| LT2 x Abony | F- | LB5000 | (Bullas and Ryu 1983) | | | unnamed intermediate | | | Spontaneous segregation of F'lac |

| Wild ancestor | Relevant marker in LB5000 | Strain name in reference | Reference | Original name (other names) | Other reference | Recipient or strain mutagenized | Donor if gene transfer | construction, comment Reference | Mode of construction, phenotype comment |
|---|---|---|---|---|---|---|---|---|---|
| LT2 x Abony | interlab transfer, new name | TR6578 | | | | | | | Roth lab from Bullas |
| LT2 x Abony | interlab transfer, new name | G251 | | | | | | | Segall lab from Roth |
| LT2 x Abony | interlab transfer, new name | STK005 | | | | | | | Raleigh lab from Segall |

§This transduction will have changed both the assignable serovar and the lysotype of the strain.
†*These strains only make sense if derived from 4274*, instead of 4247 as listed in (Bullas and Colson 1975)
