## Supplemental File S2 for "Genome archaeology of two laboratory *Salmonella enterica enterica* sv Typhimurium"

**Zaworski et al G3 Genome report Genome Archaeology**

### File S2: Analysis of 11 kb high-similarity region in the Gifsy prophages

**Summary: Long repeat regions both confound assembly and promote recombination.**

**Terminology:** The Gifsy prophages of the *S.* Typhimurium clade are designated "-1" and "-2" depending on the site of insertion into the chromosome. In LT2, Gifsy-1 is the prophage adjacent to *rseC* (STM2637) at ~56.3 min/centisomes; Gifsy-2 is adjacent *pncB* (STM1004) at ~22.5 min/centisomes (Figueroa-Bossi *et al.* 2001; Lemire *et al.* 2007). Patches of high similarity are interspersed with insertions of distinct genes, a feature called "mosaicism" (see Embedded fig1). Strains of the independently-isolated lineage ATCC14028 have similarly mosaic prophages at these sites, with an additional example, Gifsy-3, present adjacent to *icdA* (EJJ31_13865, homolog of STM1238)(Figueroa-Bossi *et al.* 2001). The ATCC14028 lineage has been used extensively in laboratory studies due to much more virulent properties in mice.

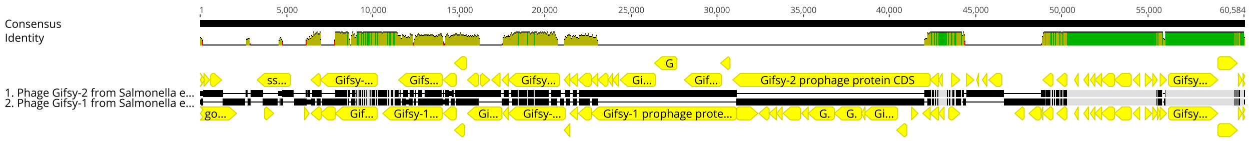


**Embedded fig1: In LT2, Gifsy-1 and Gifsy-2 are highly similar at one end:** Mauve alignment of the LT2 Gifsy prophages (extracted from NCBI: NC_003197.2 using the "source" annotation for these phages). Numbering of the consensus sums the lengths unique sequence of the two phages. Consensus Identity: green=100% identity; yellow ticks indicate various degrees of DNA identity; white spaces are nonaligned gaps. Yellow arrows: annotated CDS; black segments between CDS rows represent the DNA, while connecting lines represent gaps where unrelated genes are carried by the partner. Each phage carries an additional 4 kb of unique sequence at the end coding for distinct Int and Xis proteins not included in the alignment.

**Biology:** These phages are known to reassort by recombination and indeed share complex regulatory interactions during induction and pathogenesis (Lemire *et al.* 2007; Lemire *et al.* 2011). Long stretches of near-identical sequence could be substrates for recombination between prophages during laboratory propagation. The two Gifsy prophages of LT2 share ~11 kb of high identity (see Embedded fig1). This region codes for proteins that participate in homologous recombination, as well as regulation and replication. The recombination proteins are related to those of the Lambda recombination cluster: Red (Exo, Beta), Orf and Rap. In Lambda, Exo and Beta provide a homologous recombination system that can substitute for the host RecABCD system (MURPHY 2016). Rap (originally named NinG) facilitates recombination near linear ends via the lambda Red pathway (TARKOWSKI *et al.* 2002), acting as a structure-specific nuclease (SHARPLES *et al.* 2004).

The homologs of Lambda Exo found in Gifsy-1 (STM1009) and Gifsy-2 (STM2632) are very large (962 aa). In this, they are closer to RecE of the Rac prophage of *E. coli* K-12 than to Lambda Exo itself (LUISI-DELUCA *et al.* 1988; ZHANG *et al.* 2009). As with RecE and RecT, both Gifsy-borne LT2 homologs of Exo and Beta participate in recombination when activated by mutation in vivo (LEMIRE *et al.* 2008). Action of these phage recombination systems may promote the mosaic genomic relationships among phages that has been found (MARTINSOHN *et al.* 2008).

**Sequences and sequencing technology:** Long repeats also interfere with accurate assembly of sequence reads. Advances in sequencing technology have improved capacity to determine accurate placement of sequence variations among repeats. Genbank sequences used for this analysis are:

- LT2 sequences
    - AE006468.2 (Refseq NC_003197.2): The reference LT2 rested on reads that generally shorter than 1 kb. Restriction mapping and the use of plasmid and phage clones passed through *E. coli* enabled deconvolution of repeats (MCCLELLAND *et al.* 2001). Corrections to the original sequence were deposited Jan 3 2016 by

McClelland,M., Jain,A., Saraogi,P., Mendelson,R., Westerman,R., SanMiguel,P. and Csonka,L. (reference 3 in the NCBI sequence).

- CP014051: An independent determination of LT2 sequence, using PacBio and Illumina, assembled with HGAP3 v Nov 2014; updated sequence version deposited December 2017. Deposited by FDA database for Regulatory Grade Microbial Sequences (FDA-ARGOS). In REBASE, Org num 18099 Source ATCC cites this Genbank record. Long reads of $\geq 5$ kb are routinely achieved with PacBio. Of 62 bp that differ from NC_003197.2, all but 2 are within the region discussed in observation 1 below.

- 14028 is an independent natural isolate of *S. enterica enterica* serovar Typhimurium. Sequences of several members of the ATCC14028 lineage were described in (JARVIK *et al.* 2010), with one closed sequence determined and compared with several assemblies of other members of the lineage and with assemblies of other *Salmonella* isolates.
  - CP001363.1: ATCC14028s (JARVIK *et al.* 2010) used 454, SOLID and PCR with primer walking. Two 20 kb repeats (in Gifsy-1 and Gifsy-3) could not be accurately assembled, so the consensus of the two regions was used in both locations to close the genome. In REBASE, Org num 5627 Source T. Jarvik cites this Genbank record.
  - CP034479.1: 14028 deposited in 2018 by Key Laboratory of Microbial Technology and Bioinformatics of Zhejiang Province; sequenced 161X using Oxford Nanopore GridION, assembly with Unicycler v. Unicycler v0.4.5. In REBASE, Org num 22987 Source T. Jarvik cites this Genbank record. No publication is associated with this sequence. GridION reads can be ~10 kb (KITA *et al.* 2007).

These two deposits seem to be "the same" strain determined 10 years apart.

**File S2 Zaworski et al G3 Genome report Genome Archeology**

**STK005 Gifsy alleles:**

Three observations made during comparison of LT2 and STK005 Gifsy prophages thus may have alternative explanations: intra-lineage recombination or mis-assembly. We infer one mis-assembly and two recombination events. These observations are:

1. Introduction of a novel reading-frame-preserving 42 bp insertion and 10 SNPs into the Gifsy-1 *recET* allele of the Reference (LT2 NC_003197.2 STM1009; Embedded fig2).

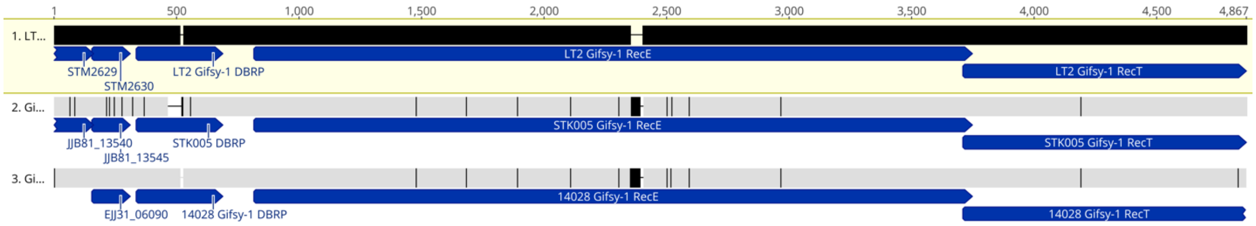


**Embedded fig2: Novel allele in STK005 is homologous to Gifsy-1 of *S* Typhimurium 14028:** *recT* and *recE* coding sequences from the Gifsy-1 phages are labelled. Row 1: LT2 reference segment; row 2: STK005; row 3: *Salmonella enterica* serovar Typhimurium 14028 CP034479.1. Locus_tags for LT2, STK005 and 14028 are respectively: RecE, Gifsy-1: STM2632, JJB81_13540, EJJ31_06080; RecT, Gifsy-1: STM2633 and JJB81_13560. Vertical lines represent discrepant positions: positions at which at least one sequence differs from the rest. Gifsy-1 of STK005 shares a significant insertion (42 nt) and 10 flanking SNPs with 14028. All of these are within the CDSs coding for homologs of RecE and RecT. At the left end of the alignment, STK005 diverges from the other two, due to a recombination patch with Gifsy-2 (see Embedded fig3; orientation is reversed from Fig 2).

This is ancestral, found in Gifsy-1 of 14028 (CP034479.1; Embedded fig2) and DT104 (Fig 3 of Lemire 2008). To explore the origin of this allele, we used PCR to ascertain the allele present in 12 LT2 isolates acquired by the Bossi laboratory, over a period of nearly 40 years, from a variety of sources (including the Salmonella Genetic Stock Center, (Calgary CA), the University of Utah and the University of Sevilla, Spain). All 12 have the "long version" of Gifsy-1 *recE* - that is, the version with 42 bp insert (see Supplementary File 4). The FDA determination of LT2 sequence agrees that the insertion and 10 flanking SNPs are located in Gifsy-1 (CP014051; not shown). This strongly suggests a mis-assembly during sequencing for original LT2 genome deposit.

2. Reciprocal exchange of sequence between Gifsy-1 and Gifsy-2 prophages.

##### 3A. streamlined view with phasing (color-coded variants)

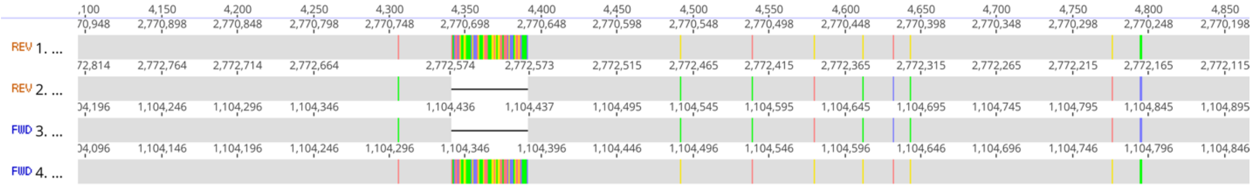


##### 3B. with CDS labels

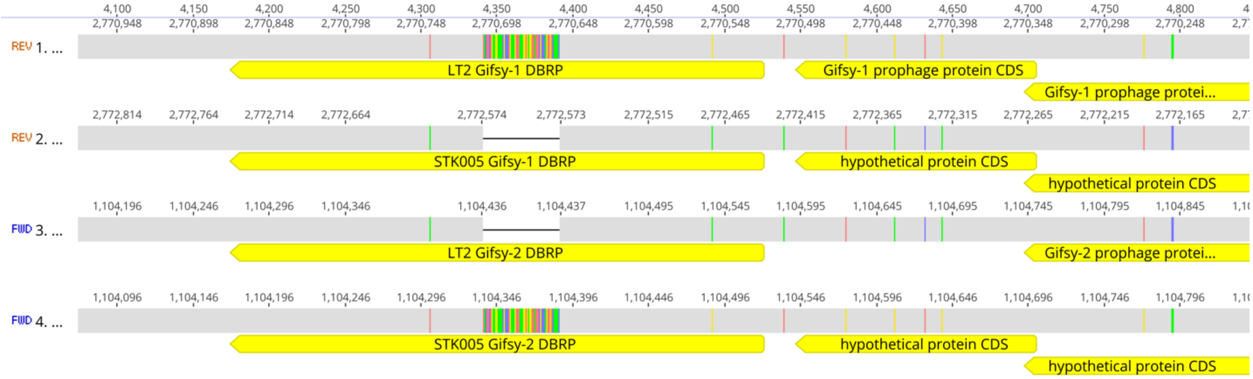


**Embedded Fig3: Reciprocal recombination between Gifsy-1 and Gifsy-2 in the descent of SKT005:** Rows: 1: LT2 Gifsy-1; 2: STK005 Gifsy-1; 3, LT2 Gifsy-2; 4, STK005 Gifsy-2.

Panel A: Streamlined view. Vertical lines represent discrepant positions, at which at least one sequence differs from the rest. At this magnification, line color is visible, enabling visualization of the phasing of variants. All 9 SNPs and the insertion are shared by STK005 Gifsy-1 (Row 2) and LT2 Gifsy-2 (Row 3); or by LT2 Gifsy-1 (Row 1) STK005 Gifsy-2 (Row 4), including a 51 bp insertion/deletion variant. No information has been lost in STK005; just moved from one prophage to another.

B. With annotations. The patch of reciprocal exchange affects three CDSs or an unannotated spacer in each phage. Recent automated annotation suggests function as a DNA breaking-rejoining protein (DBRP in the figure) for both insert and deletion alleles of STK005. The other two CDS's are of unknown function, one not even annotated in LT2 Gifsy-2. Locus tags are: LT2 Gifsy-1, (STM2631, STM2630, STM2629); STK005 Gifsy-1, (JJB81_13550, JJB81_13545, JJB81_13540); LT2 Gifsy-2, (STM1010, STM1011) and STK005 Gifsy-2, (JJB81_05180, JB81_05185, JJB81_05190).

The exchange includes a 54 bp in-frame insertion in a CDS assigned DNA breaking-rejoining activity (DRBP; Embedded fig3A, 3B). Gifsy-1 JJB81_13550 lacks the insertion; Gifsy-2 JJB81_05180 has it. In contrast, LT2 Gifsy-1 STM2631 has the insertion, while Gifsy-2 STM1010 lacks it. The FDA sequence CP014051 agrees with RefSeq LT2 (NC_003197.2) here, arguing against mis-assembly.

3. 9 SNPs in a 450 nt stretch define a gene conversion event in a Rap (NinG) homolog (Embedded fig4A, B). LT2 has distinct sequences in Gifsy-1 STM2619 and Gifsy-2 STM1021 (both sequence deposits), while STK005 has identical sequence in the two prophages (JJB81_13480). At the right, Gifsy-2 agreement resumes (LT2 unannotated; JJB81_05250).

**4A. streamlined view with phasing (color-coded variants)**

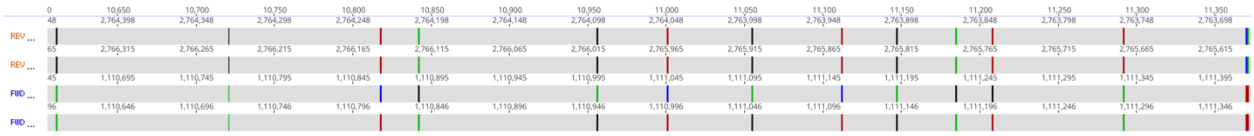

**4B. with CDS labels**

#### File S2 Zaworski et al G3 Genome report Genome Archeology

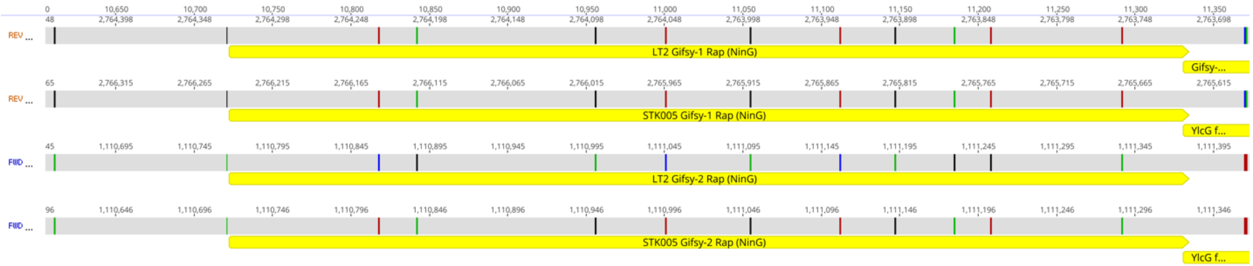


**Embedded fig4: Gene conversion: Gifsy-2 of STK005 has lost information, receiving sequence from Gifsy-1**: Rows: 1: LT2 Gifsy-1; 2: STK005 Gifsy-1; 3, LT2 Gifsy-2; 4, STK005 Gifsy-2.

Panel A: Between 10,800-11,250 of the alignment, LT2 Gifsy-2 (3rd row) carries unique sequence at 9 discrepant positions. Gifsy-2 of STK005 (row 4) agrees with Gifsy-1 of LT2 (row 1) rather than with Gifsy-2 (row 3). Unidirectional transfer of information from one allele to another (with loss of the original copy) is called gene conversion.

Panel B: All the gene-converted positions are within the gene coding for a homolog of Lambda Rap (NinG), a structure-specific nuclease.

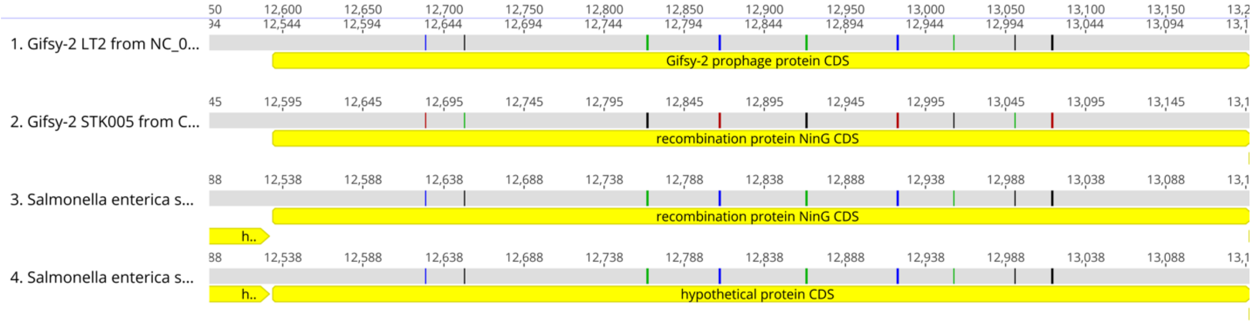
