## Supplemental File S3 for "Genome archaeology of two laboratory *Salmonella enterica enterica* sv Typhimurium"

**File S3: STK005 genotype inferences from sequence**

Describes the genotype; explains assignment of flagellar genotype; describes the protocol used to generate the variant list found in Supplementary File 1; lists protein changes that account for the genotype and some additional possible changes.

**Supplementary File 3 STK005 genotype inferences from sequence**

For sequence variants see Excel file Supplementary File 1 STK005 variants

For protocol to generate Supplementary File 1 see Section III of this document

<u>Section I</u>

Genotype of LB5000 (Salmonella Stock Center Genotype SGSC181) from Table "Supplementary Table 1 Lineage of LB5000 with classification references.docx"

*metA22 metE551 trpC2 ilv-452 H1-b H2-e,n,x (cured of Fels 2) flaA66 rpsL120 xyl-404 leu hsdL6 hsdSA29 hsdSB121*

Revised genotype from sequence and other sources (see section III):

*metA22(fs) metE551-G293R trpC2-G192D ilvC452-G140S [fliC(H1-b) (Fels 2$^-$)]Abony2 [fljB(H2-e,n,x)]Abony1 flaA66fs rpsL120-K43R xylR404-G82E leuA-D142N res6-D917N hsdSA29{STM4496-STM4492} hsdR121-R825C*

Genotype conventions:

A: Square brackets: regions originating from two transductions from *S enterica enterica* sv Abony SW803 are set off in brackets followed by Abony1 or Abony2. Abony1 was historically the first of the two. Comparisons with serovar Abony str. 0014 - NZ_CP007534.1 were used to identify transduction borders.

**File S3: STK005 genotype inferences from sequence**

B: nutritional markers in the genotype as received are given gene names here, identified by comparison with AE006468 (the sequence of *Salmonella enterica enterica* sv LT2 ATCC700720), followed by allele numbers if present, then amino-acid change following a hyphen: AA(WT)codon#AA(mut)

C: H-antigen terminology adopted as described below, section II.

D: names of restriction genes follows the annotation of LT2 (*res* for former *hsdL*, *hsdR* for former *hsdSB*). Mutations accounting for *hsdSA* appear in two genes, to be described in a subsequent paper. For now, these are listed as *hsdSA29{JJB81_22595-JJB81_22575}*

<u>Section II. H1, H2 and *fla66*</u>

A: H1 and H2, flagellin genes

H1: *fliC* is at 40 min; phase 1 flagellin (FliC=H1 locus)

H2: *fljB* is at 60 with *hin*; phase 2 flagellin (FljB = FliC homolog at H2)

Phase variation:

- *fljA* at 60 min H2 locus; phase 1 transcriptional repressor.
- When *hin* and the H2 promoter is flipped one way, FljB (H2) and FljA are expressed and FliC (H1) is repressed. When inverted, FljB (H2) is not expressed and FliC (H1) repression by FljA is lifted.

Much of the genetic work defining these properties was done with strains in the lineage leading to STK005, ie with the Abony configuration at H1 and H2.

B: *fla66*

*fla66* must be *fliF66fs.* FliF-84frameshift. -G, not in a run.

Logic:

**File S3: STK005 genotype inferences from sequence**

1. *fla66* must be one of *fliEFGH* at ~40 min near *fliC* (see reasoning in Genotype History below), not at ~60 min with [*hin-fljB* (FliC paralog) and *fljA*].

2. cluster *fliEFGH* is outside the Transduction A (Supplementary Table 1) region but adjacent to it.

3. STK005 (CP067397) aligns with Abony at 40 min region, but only *fliZ-nml-(fliC* homolog--should be *fljA*)-middle of *fliD*. The FlaA region (steps 1 and 2 below) is outside the transduction tract.

4. In alignment with LT2, STK005 has two changes to the N-terminal region of *fliF* of LT2 (frameshift by -1 at codon 84 (not in a run), and A->V at codon 158) , sufficient to explain failure to swim. No changes are found in *fliE,G* or *H*, which might also be possible. See highlight in step 3 below.

Genotype assignment and conventions history:

1. Joys (Joys and Stocker 1965) described *fla-66*

2. (Joys and Stocker 1969) described *flaB-flaD-flaA*-H1.

3. In Wilkinson (Wilkinson et al. 1972) this allele is referred to as *flaA66*

4. The *fla* genes were renamed (Sanderson and Roth 1988): *flaB* is now *fliP*; *flaD* is *fliQ*; *flaA* is *fliEFGH*; H1 is *fliC*. In Sanderson and Roth map of 1988 *flaA* (region at 40 min) was divided as part of a harmonization with *E. coli* nomenclature.

| | |
|---|---|
| flaAI: | *fliE* flagellar synthesis function unknown |
| flaAII.1 | *fliF* basal body M-ring protein |
| flaAII.2 | *fliG* (also was *motC*, *cheV*) |
| flaAII.3 | *fliH* flagellar synthesis function unknown |

Section III. Creating list of variant positions.

**File S3: STK005 genotype inferences from sequence**

Alignment of LT2 and STK005: Variant list generation

- Variants found using Geneious Prime 2021.0.3 Build 2020-12-23
- Mauve alignment of 22 Jan 21
- Three regions of the STK005 linear genome were inherited from LT2, separated by two regions inherited from Salmonella Abony SW803. Alignments with two extracted Abony sequence segments were used to determine the STK005 coordinate breakpoints below.
- S Abony H1 fla region v STK005 alignment: Alignment of 2 sequences: uvrC-fliS Salmonella enterica subsp. enterica serovar Abony str. 0014 - NZ_CP007534.1 extraction (bases 3 to 15412), Salmonella enterica subsp. enterica serovar Typhimurium - CP067397 (bases 2037077 to 2052486) [from "properties" tab in "Info" menu].
- S Abony H2 fla region v STK005 LCB1 alignment: Alignment of 2 sequences: Salmonella enterica subsp. enterica serovar Typhimurium - CP067397 (bases 2822320 to 2864091), S Abony fla region grpE to tonB NZ_CP007534 extraction (bases 3 to 41771).

| segment | from STK005 coordinate | to STK005 coordinate | source | note |
|---|---|---|---|---|
| 1 | 1 | 2,038,907 | LT2 | 1829 nt overlap seg 2; includes *fliD*, 3 mismatches |
| 2 | 2,037,078 | 2,052,486 | Abony H1 | 988 nt overlap w seg 3; includes fliD no mismatches |

**File S3: STK005 genotype inferences from sequence**

| | | | |
|---|---|---|---|
| 3 | 2,051,498 | 2,820,465 | LT2 |
| 4 | 2,822,320 | 2,864,091 | Abony H2 |
| 5 | 2,864,092 | 4,796,208 | LT2 |

For each segment, variant tables were generated as below. In addition to the positions of variants, some additional feature locations were included: rDNA, mobile elements, and "misc features", which include regulatory RNA leaders for amino acid biosythetic operons, a temperature-sensor RNA and a programmed frameshift in *dnaX.* Several variants are located in rDNA.

In Geneious Prime 2021.0.3 build 2020-12-23, aligned sequences generated with Mauve were processed as below.

In **Alignment view**

- Selected both sequences to the coordinates of the STK005 in the table above (**NOT** the coordinates of the **alignment**, which adds together all insertions such as mobile elements or repeat expansions).
- Extract as an alignment
- Give alignment a name: "LT2 vs STK005 alignment extraction 1", or 2, or 3; or Abony H1 or H2
- Set reference in Alignment View:

  Select sequence 1: LT2 or Abony

  control key while selecting sequence 1: set reference sequence

**In menu Annotate and Predict**

**File S3: STK005 genotype inferences from sequence**

- Find Variations/SNPS
  - Check Minimum Coverage: value 1
  - Check Minimum Variant Frequency: value 0.5
  - Find variants: Inside and outside CDS
  - Check Analyze effect of variants on translations: Bacterial code
  - Advanced options:
  - Check Merge adjacent variations
  - Check Record names of all contributing sequences for each variant
  - CDS Properties to Copy: gene, product, protein_id, Locus_tag, Note
- This generates a "track" of Variants **in the coordinates of the reference.**

Save

**With alignment in view**

- in "Advanced" options; Properties; chose numbering: all sequences
- in "annotation" options: Misc Feature, Mobile element, rRNA, variants. For Abony segments, which are short, also chose CDS.
- in "Annotations and Tracks" choose "pop out" to see table
- in "Columns" option,
  - Choose columns and arrange in order left to right
    - Sequence name
    - Minimum
    - Maximum
    - Min (original sequence)
    - Max (Original sequence)

        - Length (in the reference sequence)
        - Locus_tag
        - Name (this will be the new nucleotide(s) if a polymorphism)
        - Polymorphism type
        - Change (nucleotide)
        - CDS (occupied only if there is a polymorphism)
        - Protein effect
        - CDS codon number
        - Amino acid change
        - Product
        - Note
    - Save

      Export table as .csv; change to Excel.

Section IV. Additional genotype comments

- Leu$^-$ due to NG mutagenesis in isolation of SB mutation (Bullas pers. Comm. 15/04/81) to Sanderson; Bullas and Ryu 1983). Roth lab record for TR6578 notes leaky Pro-, and "TR6578 was not terribly viable in stabs".
- The *hin* gene is functional in this lineage. Our two sequenced and annotated derivatives have three disagreements with the Abony reference, but they agree with each other and LT2 at those positions; and inversion has occurred in the lineage between STK005 and ER3526.

Genotype inferences from the genome:

**File S3: STK005 genotype inferences from sequence**

- *metA22=metA22*=>MetA276-fs: STM4182 in Segment 5 *metA*: codon 276fs; 10 nt deletion -CCTGGCGCAG
- *metE551=metE551*=>MetE-G293R: STM3965 in Segment 5 *metE* G->A transition; G293R amino acid change
- *trpC2*=>TrpC-G182D: STM1725 in segment 1 *trpC* G->A transition; G182D amino acid change
- *ilv-452*=>IlvC-G140S+194 truncation: STM3909
- *ilvC*: in segment 5 G->A transition G140S amino acid change; G->A at codon 194->UGA
- *rpsL120*: in segment 5 RpsL-K43R: STM3448 *rpsL*: T->C transition; K43R amino acid change
- *xyl-404*: in segment 5 XylR-G82E: STM3662 *xylR* G-> A transition; G82E amino acid change
- *leu*: in segment 1 LeuA-D142N:
    - *leuA*: STM0113 *leuA* C->T transition, D142N
    - *leuO*: STM0115 *leuO* C->T transition at codon 143; no aa change
- *hsdL6*: in segment 1 Res-D917N*:* STM0358 (*res*) in LCB1 two G->A transitions; one D917N amino acid change
- *hsdSB121*: in segment 5 HsdR-R825C: STM4526 (*hsdR*) in LCB3 *hsdR*: G->A transition; R825C amino acid change; G->A at codon 293 no aa change
- *hsdSA29*: in segment 5 by hypothesis from genetic mapping and domain identification:
    - mutations are not in candidate DNA MTase STM4495
    - 9 aa changes in all in three CDS

**File S3: STK005 genotype inferences from sequence**

- 1 change in STM4489 (putative helicase): C->T; aa change A486V
- 7 changes in STM4492 (putative cytoplasmic protein): all G->A; aa changes P731S, P526L, T498I, A153V
- 5 changes in STM4496 (putative ATPase): three G->A, two C->T; aa changes G1207S, A1206T, A1144V, S935L

Markers in regions aligned with S. Abony:

- Fels2$^{-}$
- H1-b H2-e,n,x refer to alleles of flagellar antigen from S. Abony.

Other notable changes from a scan of the genes:

Segment 1 (LT2)

- CarA T253I (carbamoyl phosphate synthase; early step in aromatics synthesis; originally known in *E coli* as *arg+ura* because deficiency alleles were satisfied by supplement with both metabolites)
- PanB E251K (pantothenate metabolism)
- ProB P179S (proline synthesis)
- ProC A106V (proline synthesis)
- ProY G115R, G267D (permease)
- SpeG E172K (spermidine metabolism)

Segment 2 (Abony) carries *uvrC*, *uvrY* without changes

Segment 3 (LT2)

- ClpB P650L (chaperone)
- CbiD G184D (cobalamin sysnthesis)

Segment 4 (Abony) carries *grpE, recN* and *bam* in addition to flagellar stuff.

**File S3: STK005 genotype inferences from sequence**

Segment 5 (LT2)

- AtpD C435K (membrane ATPase)
- AtpG P190S (membrane ATPase)
- Crp G68S (catabolite regulation protein)
- MetL A23V (aspartokinase and homoserine dehydrogenase domain)
- MutL P418S (mismatch repair)
- MutS G782S (mismatch repair)
- NrdE L114P (ribonucleotide reductase II alpha chain)

Joys, T. M., Stocker, B. A. 1965. Complementation of non-flagellate Salmonella mutants. J Gen Microbiol. 41:47-55.

Joys, T. M., Stocker, B. A. 1969. Recombination in H1, the gene determining the flagellar antigen-i of Salmonella typhimurium; mapping of H1 and fla mutations. J Gen Microbiol. 58:267-275.

Sanderson, K. E., Roth, J. R. 1988. Linkage map of Salmonella typhimurium, edition VII. Microbiological reviews. 52:485-532.

Wilkinson, R. G., Gemski, P., Jr., Stocker, B. A. 1972. Non-smooth mutants of Salmonella typhimurium: differentiation by phage sensitivity and genetic mapping. J Gen Microbiol. 70:527-554.
