## Supplemental File S4 for "Genome archaeology of two laboratory *Salmonella enterica enterica* sv Typhimurium"

### PCR survey for 42bp insertion in isolates of *S. enterica* sv Typhimurium str LT2

Nineteen LT2 strains or derivatives listed in Embedded Table1 were tested: 12 obtained by the Figueroa-Bossi lab at different times from different sources, and 6 control strains cured by the Figueroa-Bossi lab of one of the two prophages (2 cured for Gifsy-1; 4 cured for Gifsy-2) (Embedded Fig1). PCR amplification employed primers flanking the site of the 42 bp insertion and three most proximal SNPs identified in File S2. The resulting fragment is 320 bp without insert and 362 bp with it.

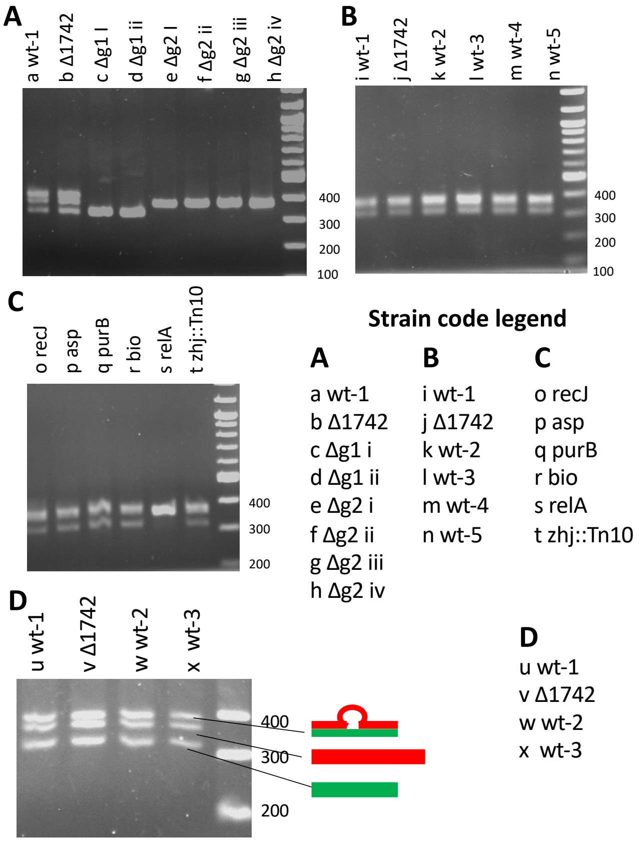


Embedded Fig1. Agarose gel electrophoresis of colony PCR diagnostic for 42 nt insertion in *recE*. Panels A-D were separate experiments. Size markers are in the unlabelled lane. Strains of Table 1 were assigned a distinguishing code: wt=wild type (5 histories); Δg1 or Δg2 are strains independently cured of one or the other prophage; 7 strains carry distinguishing genetic markers.

1. When both prophages are present two fragments are present (e.g. panel A lanes a and b). From the amplification patterns of the cured strains (Δg1 or Δg2), we deduce that the smaller fragment originates in Gifsy-2 (panel A lanes b, c) and the larger from Gifsy-1 (panel A lanes d-h).

2. The larger band is generally broader, more intense and splits into two bands upon longer gel runs (panel D). We demonstrated that a species comigrating with the top band is created when

single-species pairs (e.g. Panel A lanes d and e) are denatured and reannealed (not shown). We infer that a heteroduplex species (320:362) is formed during the PCR, and that it migrates slower than the 362 species in this gel system.

3. The insertion is likely ancestral, since all of 12 strains obtained elsewhere have the upper (362 bp) band and therefore the 42 bp insert (Panels B and C), likely in Gifsy-1.

4. Eleven strains also have the lower (320 bp) band, but one, MA3296, a *relA::Tn10 spoT1* derivative, does not show the shorter band. A deletion of all or part of the Gifsy-2 prophage could have occurred, or a unidirectional recombination/conversion-type event has transferred Gifsy-1 *recE* (with the insertion) into Gifsy-2.

5. Sanger sequencing analysis of the PCR fragments from the cured strains confirms that the larger band contains the 42 bp insert as well as the three proximal SNPs (not shown; ABI trace available on request).

A word of caution: eight strains were obtained from John Roth; the four obtained elsewhere gave an original name with a TT prefix, which usually identifies strains from Roth's lab. So, ultimately all strains used here may have a common origin.

Embedded Table1. LT2-derived Isolates Surveyed

| Strain | Genotype | Original name | Lane* | Source or reference |
|---|---|---|---|---|
| MA2285 | *Δ1742[argA - recB]* | TT17243 | b,j,v, Δ1742 | J. Roth (U of Utah) |
| MA3149 | *recJ504::MudCam* | TT16817 | o recJ | J. Casadesus (Seville, Spain) |
| MA3218 | *asp544::Tn10* | TT176 | p asp | K. Sanderson SGSC Calgary CA |
| MA3219 | *purB877::Tn10* | TT282 | q purB | K. Sanderson SGSC Calgary CA |
| MA3220 | *bio102::Tn10* | TT403 | r bio | K. Sanderson SGSC Calgary CA |
| MA3296 | *spoT1 relA21::Tn10* | TT10036 | s relA | J. Roth (U of Utah) |

| Strain | Genotype | Original name | Lane* | Source or reference |
|---|---|---|---|---|
| MA3299 | *spoT23 ts zhj1036::Tn10* | TT8980 | t zhj::Tn10 | J. Roth (U of Utah) |
| MA3408 | *Δ[Gifsy-1] i* | | c Δg1 i | (FIGUEROA-BOSSI *et al.* 1997) |
| MA3409 | *Δ[Gifsy-1] ii* | | d Δg1 ii | (FIGUEROA-BOSSI *et al.* 1997) |
| MA4409 | *Δ[Gifsy-2] zgb-8163::MudF sbcE21 zfh-8157::Tn10dTc ii* | | e Δg2 i | (FIGUEROA-BOSSI *et al.* 1997) |
| MA4585 | *Δ[Gifsy-2] sbcE21 zfh-8157::Tn10dTc ii* | | f Δg2 ii | (FIGUEROA-BOSSI *et al.* 1997) |
| MA5782 | *w.t. #1* | LT2 | a, i, u wt-1 | J. Roth (U of Utah) |
| MA6280 | *w.t. #2* | LT2 | k,w wt-2 | J. Roth (U of Utah) |
| MA6281 | *w.t. #3* | LT2 | l,x wt-3 | J. Roth (U of Utah) |
| MA6282 | *w.t.#4* | LT2 | m wt-4 | J. Roth (U of Utah) |
| MA6283 | *w.t.#5* | LT2 | n wt-5 | J. Roth (U of Utah) |
| MA7320 | *Δ[Gifsy-2] iii* | | g Δg2 iii | (LEMIRE *et al.* 2008) |
| MA7321 | *Δ[Gifsy-2] iv* | | h Δg2 iv | (LEMIRE *et al.* 2008) |

Lane: lane (a-w) of Embedded fig1 with shorthand genotype: Δ=deletion; g1, g2=Gifsy phage; i, ii, iii, iv=independent isolation of cured strain; wt-#=transfer of wild type. Some strains appear in two or three lanes; a-h, panel A; i-n, panel B; o-t , panel C; u-w, panel D.
